# CD8^+^ T-cell transcription and DNA methylation show age specific differences and lack correlation with clinical outcome in pediatric Inflammatory Bowel Disease

**DOI:** 10.1101/2020.03.30.015446

**Authors:** M Gasparetto, F Payne, K Nayak, J Kraiczy, C Glemas, Y Philip-McKenzie, A Ross, RD Edgar, D Zerbino, C Salvestrini, F Torrente, P Sarkies, R Heuschkel, M Zilbauer

## Abstract

**Background & Aims:** CD8^+^ T-cell gene expression has been implicated in the pathogenesis of Inflammatory Bowel Diseases (IBD) and has been shown to correlate with disease outcome in adult patients. Moreover, CD8^+^ T-cell exhaustion was identified as a contributing mechanism that impacts on disease behaviour. We aimed to explore CD8^+^ T-cell gene expression and DNA methylation in children newly diagnosed with IBD and test their correlation with disease outcome.

**Methods:** We prospectively recruited 112 children with IBD at the point of diagnosis and 19 non-IBD controls. Follow-up samples were obtained from a subset of patients at 3-month intervals (n=62). CD8^+^ T-cells were purified from peripheral blood samples using magnetic bead sorting and genome-wide transcriptional (n=192) and DNA methylation (n=66) profiles were generated using Affymetrix and Illumina arrays respectively. Publicly available adult CD8^+^ T-cell transcriptomes and DNA methylomes were included in data analyses to investigate age dependant differences.

**Results:** Variation amongst CD8^+^ T-cell transcriptomes obtained from children showed association with disease, systemic inflammation, age and gender but lacked correlation with disease outcome in pediatric IBD. In contrast to CD8^+^ T-cell transcriptomes in adult Crohn’s Disease (CD), samples from pediatric patients did not show variation within genes forming part of the previously reported prognostic expression or T-cell exhaustion signatures. Pediatric CD patient derived DNA methylation profiles also lacked correlation with disease outcome but in comparison to adult CD8^+^ methylomes showed a higher predicted proportion of CD8^+^ naïve T-cells.

**Conclusions:** Our findings indicate age-related differences in IBD pathogenesis and highlight the importance of validating adult clinical biomarkers in pediatric cohorts.

## Introduction

Inflammatory bowel diseases (IBD) such as Crohn’s Disease (CD) and Ulcerative Colitis (UC) are complex conditions which vary vastly in their phenotype. Amongst the most prominent factors known to impact on disease presentation and behaviour is the age of onset, as patients diagnosed in childhood tend to suffer from a much more aggressive and treatment resistant disease^1^. Although this strongly suggests that key factors involved in the pathogenesis of IBD differ according to when the disease first manifests, our understanding of specific cell types or molecular mechanisms involved remains limited. CD8^+^ T-cells have been implicated in the pathogenesis of several immune mediated diseases including IBD^2, 3^. An important finding linking CD8^+^ T-cell biology to the pathogenesis of immune mediated diseases was the discovery of a prognostic transcriptional signature in adult patients diagnosed with several conditions including Systemic Lupus Erythematosus (SLE), Anca Associated Vasculitis (AAV) as well as IBD^2, 4^. Specifically, expression of distinct signature genes was found to divide patients into two groups which differed in their disease behaviour. Moreover, T-cell exhaustion was proposed as an underlying mechanism as patients displaying an ‘exhausted’ expression signature were shown to suffer from a milder disease course^5^. In addition to gene transcription, increasing evidence points towards a major role for epigenetic mechanisms such as DNA methylation in regulating fundamental aspects of CD8^+^ T-cell function, including proliferation, activation, and T-cell exhaustion^6, 7^.

Given that existing evidence in this area has been exclusively derived from studies performed in adult IBD populations combined with a large body of evidence supporting age related differences in T-cell function, we set out to investigate CD8^+^ T-cell biology in childhood onset IBD. We prospectively recruited a cohort of 131 children newly diagnosed (treatment naïve) with IBD (n=112) and non-IBD controls (n=19, Table 1) and isolated CD8^+^ T-cells from peripheral blood samples. All patients were followed-up for a minimum of 18 months from diagnosis and detailed clinical phenotype as well as outcome and treatment data were documented (Supplementary Table 1). Additionally, longitudinal samples were obtained from a subset of patients at 3-month intervals (n=62 samples). In total, we generated 193 CD8^+^ T-cell transcriptomes and 66 DNA methylomes and set out to analyse these datasets with a view to identify variation amongst patients and potential correlation with clinical outcome. Importantly, the use of publicly available datasets allowed us to test for the presence of age-of-onset specific differences in CD8^+^ T-cell molecular profiles.

**Table 1:**
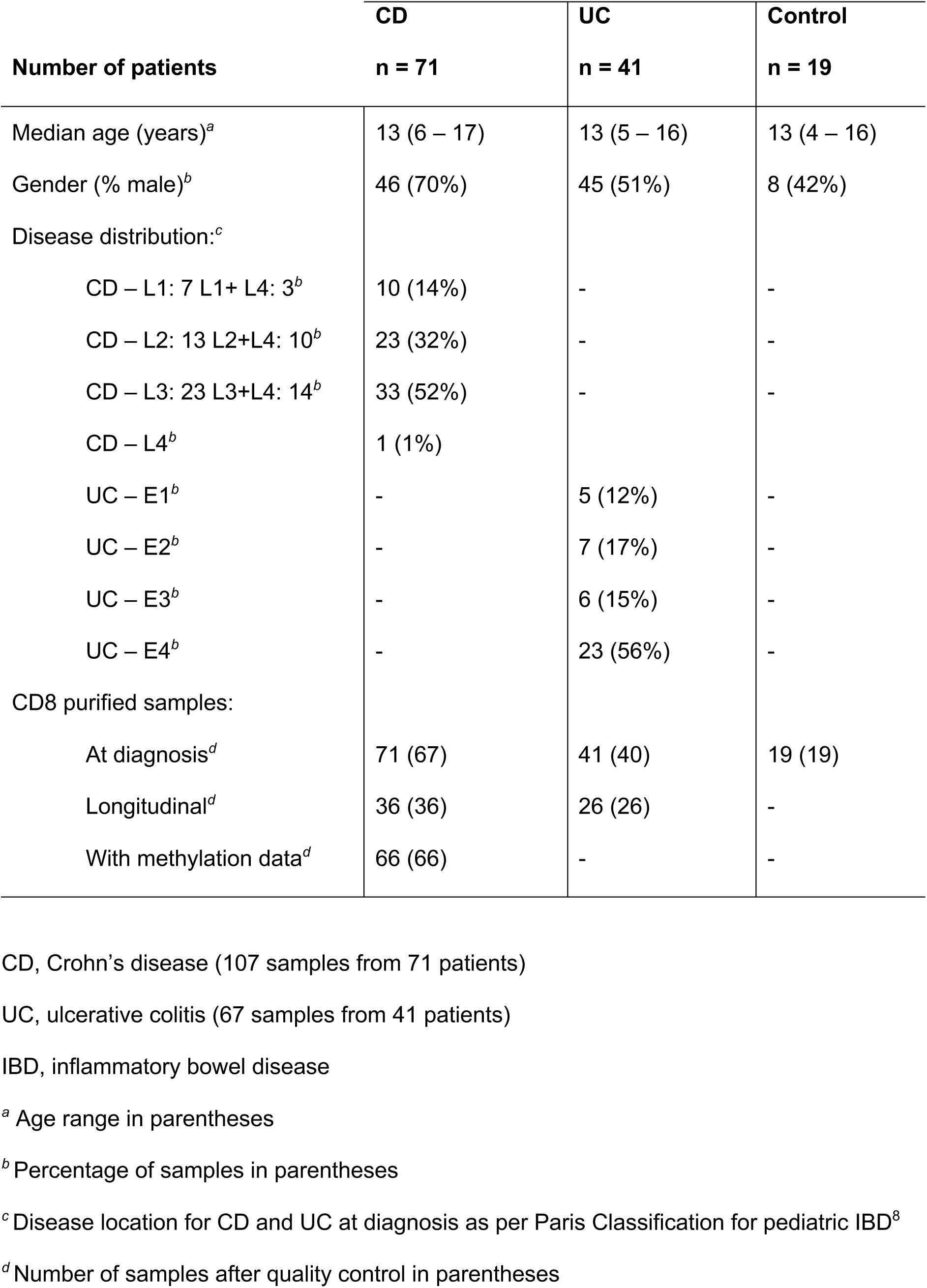
Summary of patients, samples and molecular profiles.

|  | **CD** | **UC** | **Control** |
| --- | --- | --- | --- |
| **Number of patients** | **n = 71** | **n = 41** | **n = 19** |
| Median age (years)<sup>a</sup> | 13 (6 – 17) | 13 (5 – 16) | 13 (4 – 16) |
| Gender (% male)<sup>b</sup> | 46 (70%) | 45 (51%) | 8 (42%) |
| Disease distribution:<sup>c</sup> |  |  |  |
| CD – L1: 7 L1+ L4: 3<sup>b</sup> | 10 (14%) | - | - |
| CD – L2: 13 L2+L4: 10<sup>b</sup> | 23 (32%) | - | - |
| CD – L3: 23 L3+L4: 14<sup>b</sup> | 33 (52%) | - | - |
| CD – L4<sup>b</sup> | 1 (1%) |  |  |
| UC – E1<sup>b</sup> | - | 5 (12%) | - |
| UC – E2<sup>b</sup> | - | 7 (17%) | - |
| UC – E3<sup>b</sup> | - | 6 (15%) | - |
| UC – E4<sup>b</sup> | - | 23 (56%) | - |
| CD8 purified samples: |  |  |  |
| At diagnosis<sup>d</sup> | 71 (67) | 41 (40) | 19 (19) |
| Longitudinal<sup>d</sup> | 36 (36) | 26 (26) | - |
| With methylation data<sup>d</sup> | 66 (66) | - | - |

CD, Crohn's disease (107 samples from 71 patients)
UC, ulcerative colitis (67 samples from 41 patients)
IBD, inflammatory bowel disease
<sup>a</sup> Age range in parentheses
<sup>b</sup> Percentage of samples in parentheses
<sup>c</sup> Disease location for CD and UC at diagnosis as per Paris Classification for pediatric IBD<sup>8</sup>
<sup>d</sup> Number of samples after quality control in parentheses

## Methods

### Ethical approval

Ethical approval was obtained from the local research committee (REC 12/EE/0482 and REC 17/EE/0265) and patients were prospectively recruited following informed consent. All investigations were carried out according to the Declaration of Helsinki and Good Clinical Practice Guidelines.

### Patient recruitment and clinical data recording

A total of 131 children aged between 4 and 17 (median 13) years were recruited prospectively between March 2013 and March 2016 at Cambridge University Hospitals NHS Foundation Trust in the Department of Paediatric Gastroenterology, Hepatology and Nutrition. Diagnosis of Inflammatory Bowel Disease (IBD) was made according to current international guidelines (revised Porto criteria)^8^ and included measurement of serum and stool inflammatory markers, upper and lower gastrointestinal endoscopies, radiological and histological examination. Any patient with gastrointestinal and/or extra-intestinal diseases other than IBD was excluded from the study. In total, 71 patients were diagnosed with CD, 41 with UC and 19 were classed as non-IBD, healthy controls. The latter were defined as patients who underwent endoscopic examination as part of their routine clinical care and were found to have normal macroscopic and histological appearance for their intestinal mucosa and complete resolution of any gastrointestinal symptoms.

All patients were followed up for a minimum of 18 months in the Cambridge pediatric gastroenterology unit, and detailed clinical data was prospectively recorded using the hospital’s patient electronic database (EPIC). This included phenotypic parameters at diagnosis (e.g. presence of diarrhoea, rectal bleeding, weight loss, extra-intestinal manifestations, perianal disease); disease activity scores: Pediatric Crohn’s Disease Activity Index (PCDAI)^9^ and Pediatric Ulcerative Colitis Activity Index (PUCAI)^10^; and information on disease course and outcomes. The latter covered number of treatment escalations, treatment history, response to treatment, requirement for surgical intervention, treatment with biologics and the number of unplanned inpatient admission days (Supplementary Table 1). In order to account for the fact that disease outcome is not restricted to a single measure, we designed a severity score which considered the number of treatment escalations, escalation to treatment with biologics, peri-anal disease, surgery as well as unplanned/urgent inpatient admissions (Supplementary Table 2). The score was calculated at 18 months from diagnosis and patients were categorised into mild, moderate and severe groups following blinded scoring by at least 2 consultant pediatric gastroenterologists.

### Blood sampling and processing

A peripheral blood sample (volume varied according to age from 10 to 25 ml) for the purification of CD8^+^ T-cells was taken on the same day as diagnostic endoscopy was performed. All children were treatment naïve at this time point. Additional longitudinal samples (n=62) were taken from a subset of children at 3-month intervals post diagnosis. Disease status was recorded as either ‘in remission’ or ‘active disease’ based on their clinical and biochemical disease activity scores PCDAI^9^ and PUCAI^10^ and the treatment history recorded (Supplementary Table 1). All samples were processed immediately and CD8^+^ T-cells isolated using magnetic bead sorting as detailed below.

### Magnetic bead sorting

A peripheral blood sample of 10 ml (age 4 – 10 years) or 25 ml (age 10 – 17 years) was obtained and CD8^+^ T-cells extracted using magnetic bead sorting. Briefly, peripheral blood mononuclear cells (PBMCs) were isolated by density centrifugation over Ficoll (Histopaque 1077) and CD8^+^ T-cells were separated by magnetic cell sorting using anti CD8 microbeads (Miltenyi Biotech) as described by the manufacturer. Separation was performed on an AutoMACs Pro Separator (Miltenyi Biotech). Cell purity was regularly assessed on a subset of random samples using Flow cytometry (see Supplementary Methods). With an average of 84 % (72.5 - 93.8 %), purity of our samples was found to be similar or higher compared with other published datasets.

### DNA and RNA extraction

DNA and RNA were extracted simultaneously from isolated CD8^+^ T-cell samples using AllPrep MiniKit (Qiagen), according to the manufacturer’s instructions. DNA and RNA quality were assessed using an Agilent Bioanalyser 2100 and quantified by spectrophotometry using a NanoDrop ND-1000 spectrophotometer. DNA was bisulfite-converted using Zymo DNA methylation Gold kit (Zymo Research).

### Genome wide transcriptional and DNA methylation profiling

Whole genome transcript analysis was performed on 200 ng of total RNA using the Affymetrix Human Gene ST version 2.0 Array (Affymetrix). Genome-wide DNA methylation was profiled using the Illumina EPIC platform (Illumina, Cambridge, UK). All microarray data have been deposited in ArrayExpress, accession numbers: E-MTAB-7923 (expression data) and XXXX (methylation data). DNA was bisulfite-converted using Zymo DNA methylation Gold kit (Zymo Research).

### Publicly available datasets

Gene expression microarray data (Affymetrix Human Gene ST v1.0 and 1.1) from a previously published adult patient cohort diagnosed with CD (n=35) and UC (n=32) was downloaded from ArrayExpress (E-MTAB-331)^4^. Clinical phenotype data available for this cohort included age, gender and disease severity classed as either severe (termed IBD1) or mild (termed IBD2). Secondly, we obtained publicly available CD8^+^ T-cell derived genome wide DNA methylation profiles (K450 Illumina DNA methylation arrays) from two previously published adult patient cohort. ‘Adult cohort 1’ (GSE87640)^11^, contained a total of 56 CD8^+^ T-cell samples; 18 CD, 19 UC and 19 healthy control individuals (age range 18 to 63 years). ‘Adult cohort 2’ included samples obtained from a younger (age 22–34, n=50) and older (age 73– 84, n=50) group of healthy adults (GSE59065)^12^.

### Bioinformatic analyses

A detailed description of bioinformatic methods used in this publication is provided in the supplemental methods section. Briefly, all analyses were performed in R version 3.5.2. Pre-processing of Affymetrix gene expression array data included normalization of raw signal intensity data using the variance stabilization and calibration with robust multi-array average (VSNRMA) method as part of the *affy* package *(v1*.*56*.*0)*^*13*^, and quality control assessment using *arrayQualityMetrics (v3*.*34*.*0)*^*14*^. Samples failing quality control were removed, and batch correction was performed using the “ComBat” function as part of the *sva* package (*v3*.*26*.*0*)^15^. Data was annotated using the *hugene20sttranscriptcluster*.*db* package. A total of 67 CD, 40 UC, 19 control and 62 follow-up pediatric patient samples were retained for downstream analysis. WGCNA analyses were performed on normalised and batch corrected datasets using the *WGCNA* package (*v1*.*63*)^16^ and resulting modules were correlated with clinical phenotypes as described previously^17^. DNA methylation data was processed using the *minfi* package (*v*.*1*.*28*.*0*)^18^ and included functional normalization^19^ and quality control assessment as described previously^20^. Published datasets included in this study were subjected to the same analyses. Epigenetic age and T-cell abundances were calculated using an established method developed by Horvath^21^.

## Results

### Variation of CD8^+^ T-cell gene transcription shows association with disease, age, systemic inflammation and gender

Transcriptional plasticity of CD8^+^ T-cells occurs during systemic inflammation and distinct differences have been reported in patients diagnosed with chronic inflammatory conditions including IBD^22, 23^. In order to determine the degree of variation in CD8^+^ T-cell gene transcription within our sample cohort, we first performed principal component analyses (PCA) and tested the correlation between observed variance and phenotype at diagnosis. For these analyses we included samples passing quality control obtained from children at the point of diagnosis (treatment naïve, UC n=40, CD n=67, non-IBD controls n=19). Variation in CD8^+^ T-cell gene transcription was found to be significantly associated with diagnosis (i.e. difference between IBD and non-IBD controls; Figure 1A) and age (Supplementary Figure 1A), both contributing towards the largest proportion of variance observed in PC1. Indeed, although a major overlap between IBD and control patient derived samples was observed with 89% (i.e. 17 out of 19) of control samples clustering closely together, a proportion of IBD samples separated in PC1 (Figure 1B). Additional associations were identified for disease distribution, disease subtype (i.e. UC versus CD) as well as albumin and a number of serum inflammatory markers such as platelet count, Erythrocyte Sedimentation Rate (ESR) and C-Reactive Protein (CRP). Furthermore, gender was found to be strongly associated with variance observed in PCs 8 −10 (Figure 1A, Supplementary Figure 1B). In order to investigate potential transcriptional changes over time and in response to treatment, we obtained longitudinal blood samples (n=62) and isolated CD8^+^ T-cells from a subset of patients at various stages post-diagnosis including during early remission (3 months post induction), sustained remission (6 months post induction), first and second relapse (Supplementary Table 1). Although we did not observe major differences in CD8^+^ gene expression based on specific treatment received (data not shown), samples obtained from patients in clinical remission appeared to cluster more closely to non-IBD controls (Figure 1C), further suggesting an impact of systemic inflammation on CD8^+^ T-cell transcription.

**Figure 1:**
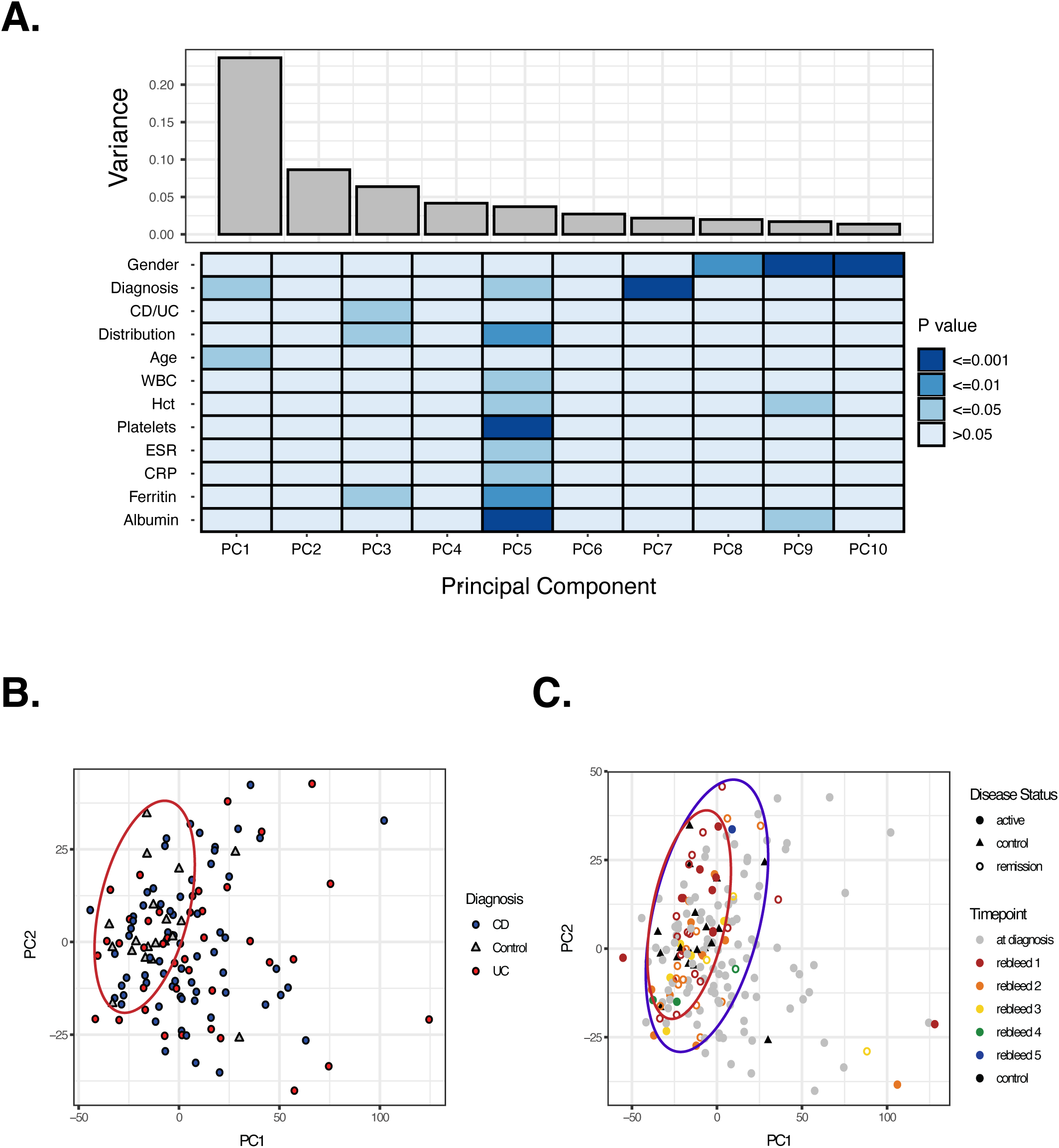
Genome wide transcriptional profiles of CD8^+^ T-cells obtained from children newly diagnosed with Crohn’s Disease (n=67), Ulcerative Colitis (n=40) and healthy controls (n=19). Samples were obtained at diagnosis (A&B, n=126) as well as during follow up (n=62, included in C). **A)** Observed variance within CD8^+^ T-cell transcriptomes (top panel) in each principle component (PC). Heat map displaying correlation between observed transcriptional variance and phenotype as well as selected serum markers at diagnosis (bottom panel). P values were generated with a Kendall correlation for continuous variables, or an ANOVA for categorical. **B)** PCA plot of CD8^+^ T-cell transcriptomes illustrating close clustering of samples derived from non-IBD controls (red circle containing 17/19 control samples). **C)** PCA plot of samples obtained from patients at diagnosis and during follow up, illustrating overlap between the majority of samples derived from patients in clinical remission (blue circle, 30/32) and non-IBD controls (red circle 17/19).

In summary, variation observed in CD8^+^ T-cell transcriptional profiles obtained from children with IBD and controls showed significant association with diagnosis, age, systemic inflammation and gender.

### CD8^+^ T-cell transcription in childhood IBD lacks correlation with disease outcome

CD8^+^ T-cell gene transcription has been shown to correlate with disease outcome in adult patients with several immune mediated diseases including IBD^2, 4^. Using publicly available adult CD patient derived CD8^+^ transcriptomes, we reproduced previously reported analyses and generated the prognostic expression signature (Supplementary methods)^4^. As shown in Figure 2A, unsupervised clustering of these samples according to expression of signature genes separated patients into two distinct groups that have been reported to vary in their disease outcome (termed IBD1=severe; IBD2=mild, based on number of treatment escalations). We next applied the same prognostic expression signature to our pediatric dataset, selecting genes comprising the adult signature and subjecting the resulting expression profiles to unsupervised clustering (Supplementary methods). As shown in Figure 2B, in contrast to the adult dataset, CD8^+^ T-cell expression of signature genes in pediatric CD (n=67) did not lead to any significant clustering. Similar results were obtained for UC patient derived samples (n=40, Figure 2C), and when combining all samples, including CD, UC, non-IBD controls as well as longitudinal samples (n=188, Figure 2D).

**Figure 2:**
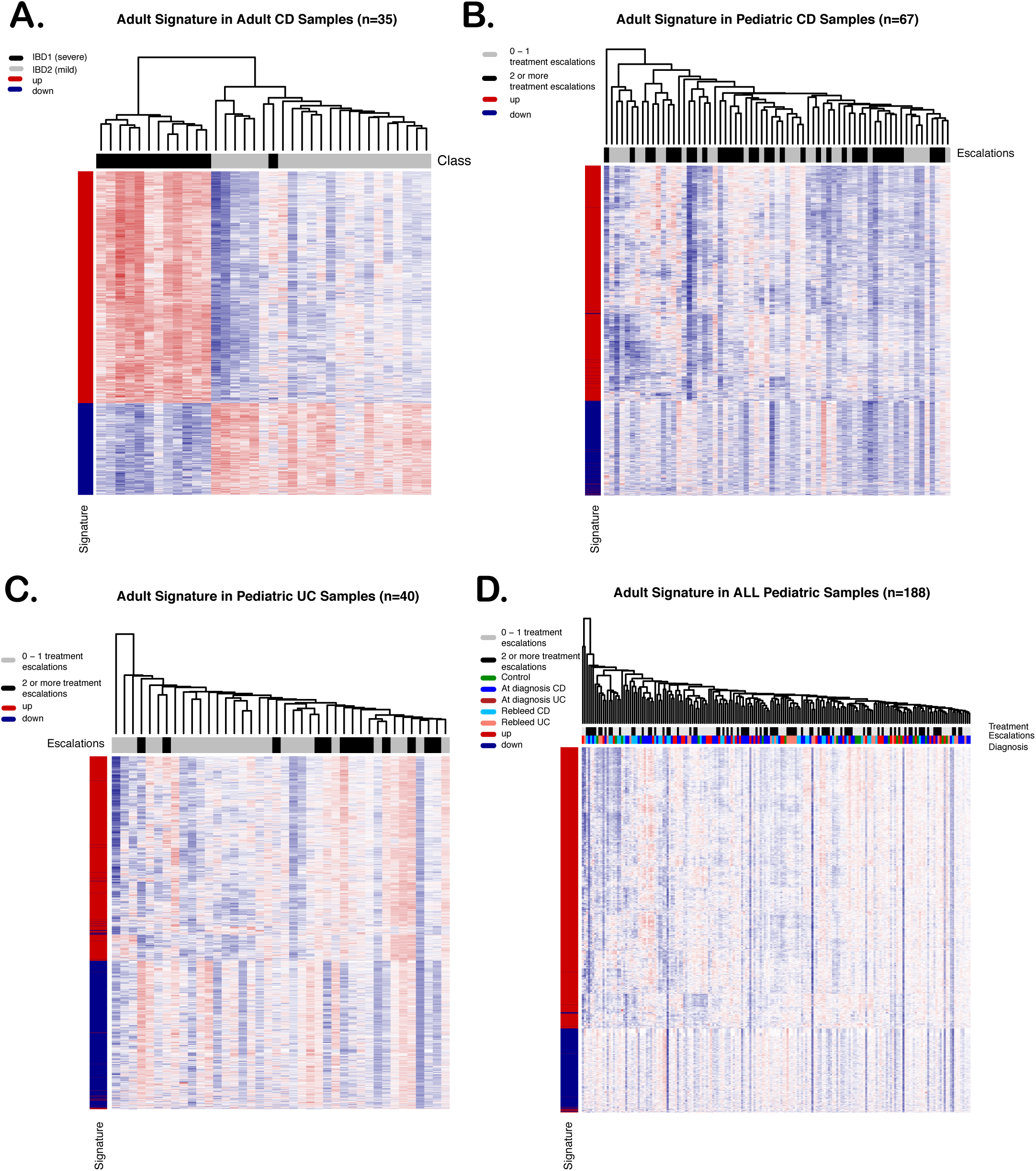
Disease prognostic transcriptional signature in adult and pediatric CD8^+^ T-cell samples. Genes forming the transcriptional signature were selected from genome-wide transcriptomes and subjected to hierarchical clustering. **A)** Heat map and hierarchical clustering of genes forming the IBD prognostic expression signature in CD8^+^ T-cells obtained from adult patients diagnosed with CD (n=35). **B-D)** Heat map and hierarchical clustering tree for genes forming part of the adult IBD prognostic expression signature applied to CD8^+^ T-cell transcriptomes of children newly diagnosed with CD (n=67, B), UC (n=40, C) and all generated pediatric transcriptional profiles (n=188, D). Clustering was tested for statistical significance using M3C.

Given the absence of the previously reported adult expression signature, we next tested our entire dataset for the presence of a potential childhood onset specific prognostic signature by correlating expression data with clinical outcome. Amongst other strategies we performed weighted gene co-expression network analyses (WGCNA) allowing the identification of gene groups (called Eigengenes or modules) that correlate with clinical outcome measures as described previously (Supplementary Methods)^10^. Applying WGCNA to adult CD patient derived expression data revealed several strong gene expression modules, some of which significantly correlated with reported patient outcome status (i.e. IBD1/IBD2, Figure 3Ai). A major overlap of the genes forming these modules with the previously reported prognostic expression signature was observed (Figure 3Aii). Moreover, expression of genes defining the top module was found to cluster patients according to outcome status confirming the validity of this analytical approach (Figure 3Aiii). In contrast to results obtained from the adult CD dataset, none of the gene modules resulting from the analysis of the pediatric CD dataset correlated significantly with any of the clinical outcome parameters tested, including number of treatment escalations, requirement for treatment with biologics, surgery, or with a summary disease outcome score designed to account for the fact that disease outcome is not defined by a single measurement (see Supplementary Methods for details; Figure 3Bi). The validity of our approach in the pediatric sample cohort was confirmed by the identification of module-trait relationships for gender and age, although the latter only reached statistical significance in UC patient derived samples (Figure 3Bii and Supplementary Figure 2).

**Figure 3:**
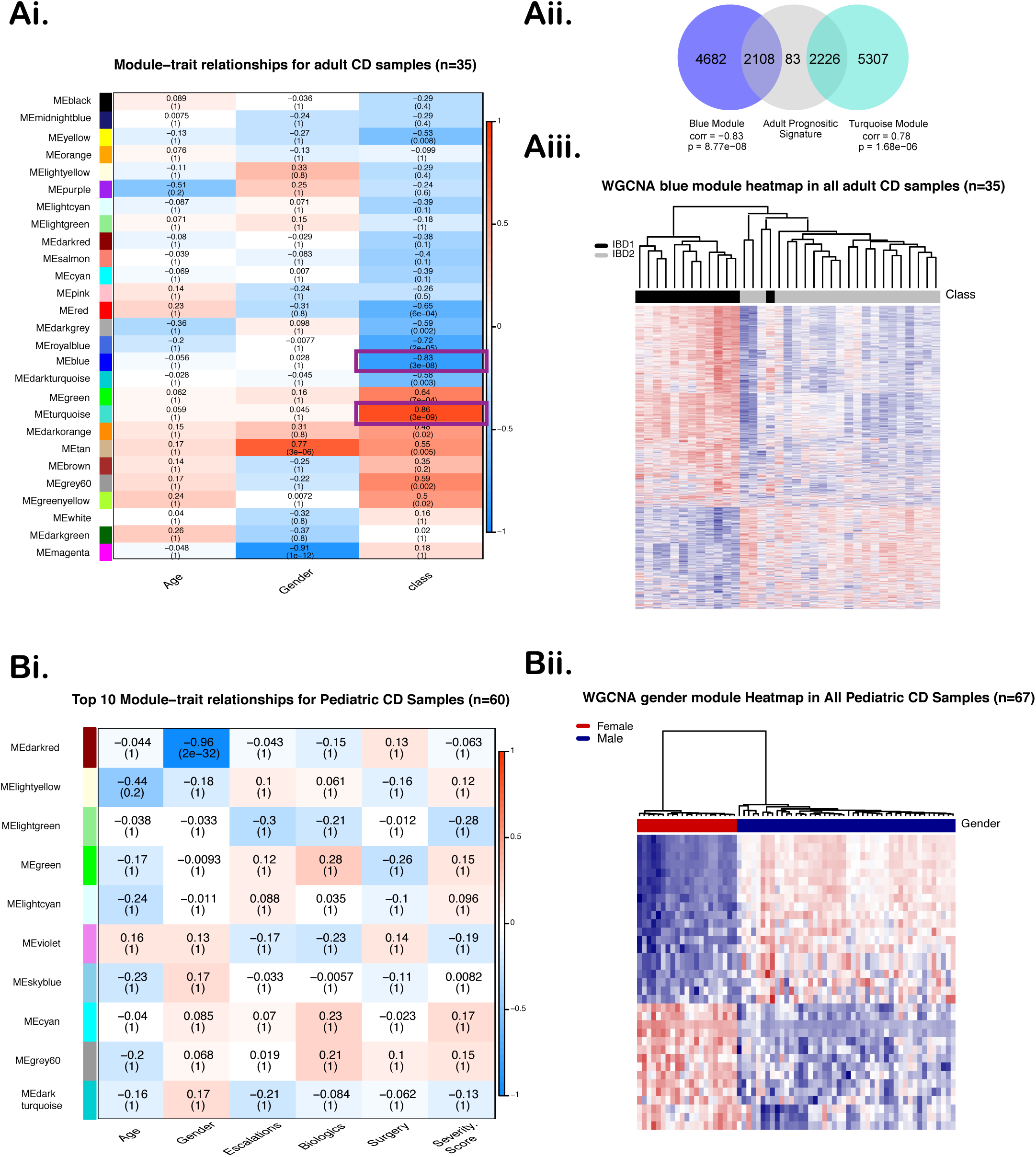
Weighted gene co-expression network analyses (WGCNA) of CD8^+^ T-cell transcriptomes derived from adult **(A)** and childhood **(B)** onset Crohn’s Disease. **Ai)** Module-trait relationship heatmap displaying gene modules and their correlation with age, gender and outcome class (severe versus mild). Numbers indicate degree of correlation between module and trait (top) and corrected p-value evaluating statistical significance of association (in paretheses). **Aii)** Overlap between genes forming the adult IBD prognostic expression signature derived from Figure 1A (grey) and top correlated modules (Blue and Turquoise). **Aiii)** Hierarchical clustering and heatmap of genes forming the Blue module. **Bi)** Module-trait relationship heatmap of CD8^+^ T-cells derived from children newly diagnosed with CD (n=60). **Bii)** Hierarchical clustering and heatmap based on genes forming the gene module associated with gender (Darkred). Samples of patients commenced on treatment with biologics at diagnosis were excluded from these analyses.

Taken together, our results demonstrate a lack of correlation between CD8^+^ T-cell transcription and disease outcome parameters in children newly diagnosed with IBD and highlight distinct differences to previously published datasets derived from adult IBD patients.

### Expression of genes associated with CD8^+^ T-cell exhaustion show limited variation in pediatric patient samples

T-cell exhaustion has been reported as an underlying mechanism contributing to the observed differences in adult onset IBD disease behaviour^5^. To investigate these findings in our patient cohort, we performed unsupervised clustering analyses based on genes associated with T-cell exhaustion as reported previously^5^. Whilst expression of exhaustion related genes^24^ clustered adult CD patient derived transcriptomes into distinct sub-groups (Figure 4A), we did not observe any significant clustering in CD8^+^ T-cell transcriptomes of children newly diagnosed with CD, UC or when combining all pediatric samples (n=188, Figure 4B and Supplementary Figure 3).

**Figure 4:**
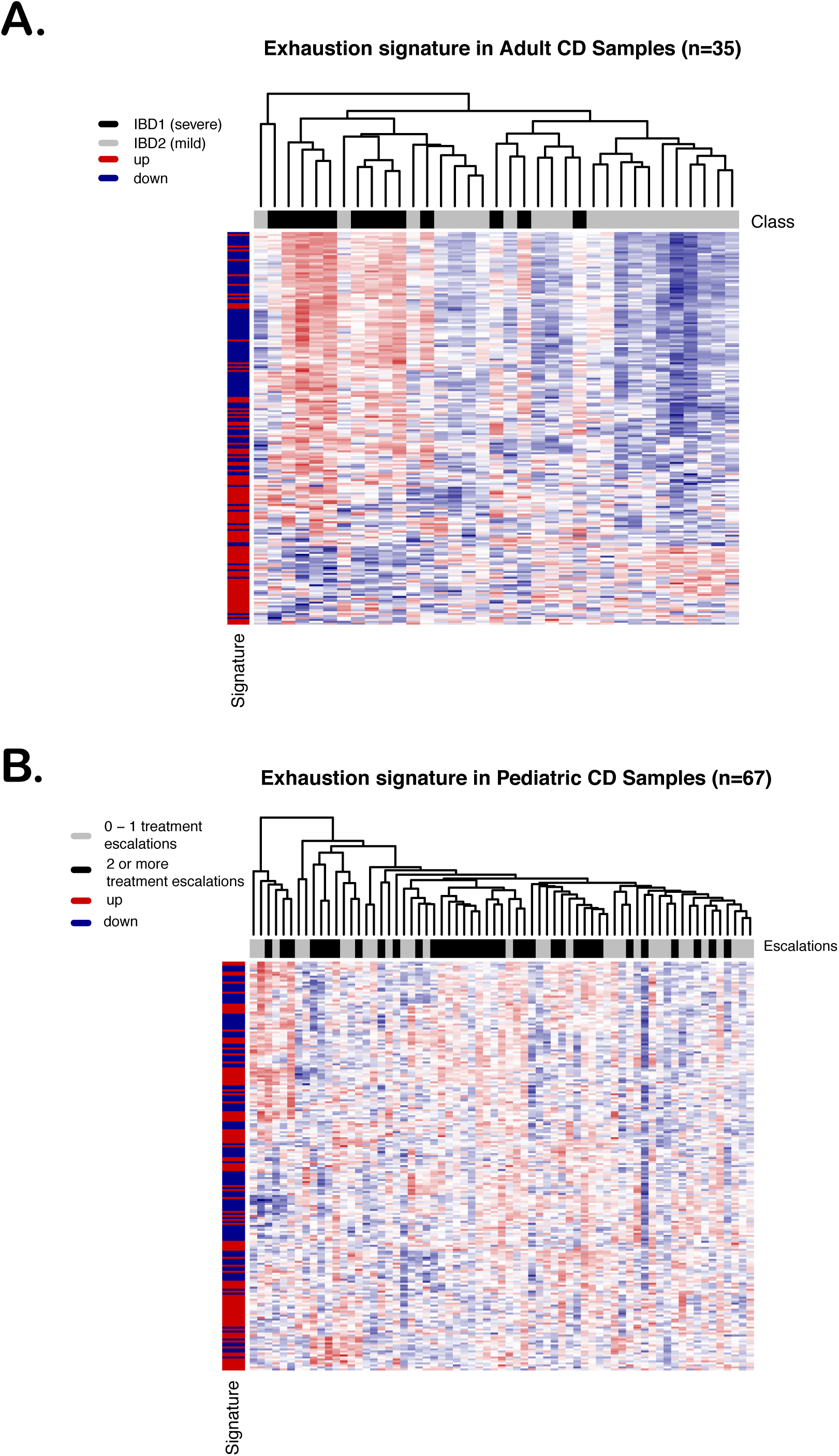
Transcriptional variation of genes associated with T-cell exhaustion in CD8^+^ T-cells. Genes forming a transcriptional T-cell exhaustion signature^24^ were selected from genome-wide transcriptomes and subjected to hierarchical clustering. Heatmap and hierarchical clustering trees are displayed for samples obtained from adults (**A**, n=35) and children (**B**, n=67) diagnosed with Crohn’s Disease.

Together these findings suggest that T-cell exhaustion associated gene transcription does not vary significantly in CD8^+^ T-cells obtained from children diagnosed with IBD or non-IBD controls.

### CD8^+^ T-cell DNA methylation shows association with age but not with disease outcome in children newly diagnosed with CD

DNA methylation has been shown to regulate key aspects of CD8^+^ T-cell function, including cellular differentiation and exhaustion^7^. Moreover, epigenetically mediated alterations in T-cell function have been implicated in disease pathogenesis of several immune mediated conditions^6, 25, 26^. We performed genome-wide DNA methylation profiling of CD8^+^ T-cells derived from children newly diagnosed with CD (n=66). Variation observed amongst these samples showed significant associations with disease distribution, gender, age and inflammatory markers (Figure 5A). Similar to CD8^+^ T-cell transcriptomes, we did not detect any significant correlation between DNA methylation and clinical outcome using various analytical approaches including differential methylation and variance decomposition analyses (Figure 5B and data not shown). In order to investigate potential age-related changes in CD8^+^ T-cell DNA methylation, we included publicly available genome wide methylomes obtained from 2 adult cohorts^11, 12^, one of which included samples obtained from adult IBD patients (Adult Cohort 1)^11^. Firstly we applied an ‘epigenetic clock’ algorithm^21^ to these datasets and confirmed a significant correlation between the predicted age based on DNA methylation and chronological age (p<0.005; Figure 5C). As DNA methylation has been used as a highly accurate measure to predict cellular composition, we next estimated the proportion of naïve and memory CD8^+^ T-cells in our dataset as well as the published adult datasets. As shown in Figure 5D, in both adult datasets we observed a significant inverse correlation for the estimated proportion of naïve CD8+ T-cells with age as well as an age-dependant increase in the proportion of memory CD8^+^ T-cells in Adult cohort 2 (p<0.005). Interestingly, amongst the CpGs which were found to display age dependant methylation changes, 265 were located near genes implicated in T-cell exhaustion and/or cell differentiation such as CD160 and Programmed Cell Death 1 (PDCD1, Supplementary Table 3, Supplementary Figure 4).

**Figure 5:**
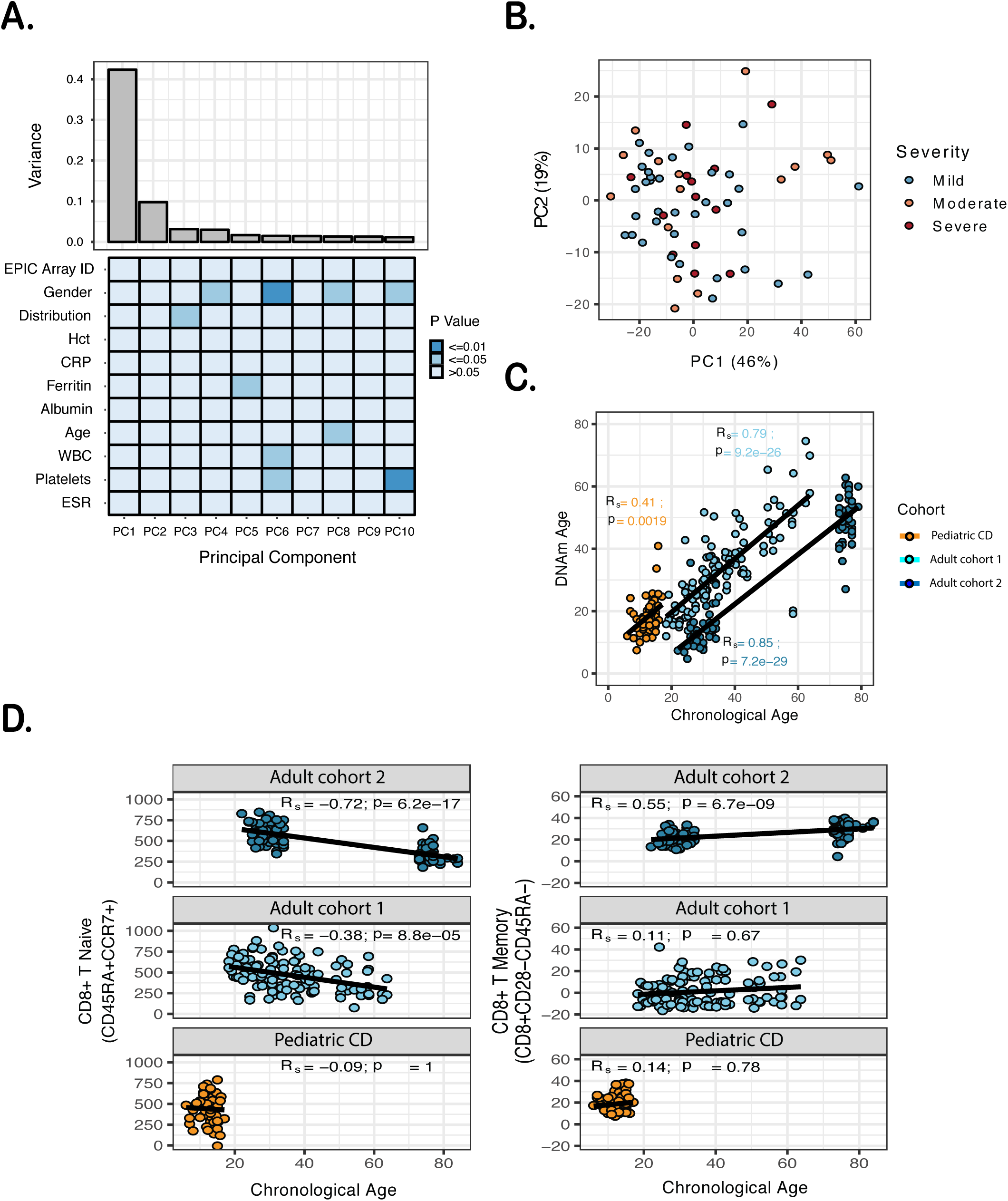
Genome-wide DNA methylation profiles derived from CD8^+^ T-cells. **A)** Variation observed in CD8^+^ T-cell DNA methylomes obtained from children newly diagnosed with Crohn’s Disease (n=66). Degree of variance in each principal component (PC – top panel) and association with phenotype/serum parameters are indicated (heatmap, bottom panel). P values were generated with a Spearman correlation for continuous variables or an ANOVA for categorical. **B)** PCA plot of childhood CD patient derived CD8^+^ T-cell DNA methylomes. Samples labelled according to disease outcome (mild, moderate, severe). **C)** ‘Epigenetic clock’ applied to CD8^+^ T-cell DNA methylomes derived from children newly diagnosed with CD as well as two publicly available datasets (Adult cohorts 1 and 2). Adult cohort 1 includes samples derived from adult patients diagnosed with IBD and healthy controls^11^. **D)** Estimated proportion of naïve (left) and memory (right) CD8^+^ T-cells based on DNA methylomes demonstrating correlation with age in adult cohorts. Adult cohort 1 (n=56)^11^, Adult cohort 2 (n=100)^12^

Together, these results further support the lack of CD8^+^ T-cell derived molecular profiles with disease outcome in children newly diagnosed with CD and highlight age as a likely factor impacting on CD8^+^ T-cell function and accounting for distinct differences between patient cohorts.

## Discussion

Accurate disease prognostic biomarkers in IBD remain an essential requirement for effective personalised treatment, by predicting clinical outcomes in individual patients. However, the search for parameters that reliably predict disease behaviour at diagnosis remains challenging, with most studies identifying markers of short-term disease activity rather than longer-term clinical outcomes. An innovative and elegant approach of combining purification of individual leukocyte subsets with genome-wide transcriptional profiling led to the discovery of a prognostic expression signature in CD8^+^ T-cells and, more recently, the development of a whole blood based clinical biomarker in adult IBD patients^27^. Interestingly, the reported prognostic expression signature was also found to be present in healthy adult control cohorts^2^, strongly suggesting differences in CD8^+^ T-cell function are unrelated to chronic inflammation. Indeed, T-cell exhaustion was later identified to contribute to the observed differences. Specifically, these findings suggested that individuals displaying exhausted CD8^+^ T-cell expression phenotypes experience a milder immune mediated disease course, whilst being more susceptible to infectious diseases^5^.

Here, we investigated CD8^+^ T-cell transcription in a large, prospectively recruited cohort of children, newly diagnosed with IBD. Our findings stand in clear contrast to those described in adult patient populations, as we were unable to replicate the previously reported CD8^+^ T-cell transcriptional signature in any of our 193 genome wide profiles. Moreover, we were unable to find any correlation between CD8^+^ T-cell gene transcription or DNA methylation and clinical outcomes. This was despite including a wide range of clinically meaningful pediatric outcome measures, those used in previous adult studies (e.g. number of treatment escalations), as well as a new summary outcome score, in which we combined key aspects of disease severity. Defining disease outcome is complex. There continues to be a lack of consensus on outcomes in the search for clinical biomarkers in IBD, as described recently by Dulai and colleagues^28^. Nevertheless, despite the presence of variation in disease outcome measures within our patient cohort, the clear lack of correlation with variation of CD8^+^ T-cell gene expression or DNA methylation does not support their use as a prognostic biomarker in childhood onset IBD. Given the near-identical methodology, and the application of additional novel analytical approaches to both adult and pediatric datasets, the differences we observe are most likely to be due to the age difference between cohorts. Indeed, human ageing is well known to have a major effect on immunocompetence, generally leading to a gradual decline^29^. Specifically, CD8^+^ T-cell populations have been reported to be particularly affected by aging, as their ability to generate primary CD8^+^ T-cell responses to newly encountered antigens decreases over time^30^. This effect can be partly attributed to a decrease in the number of naïve CD8^+^ T cells with advancing age, as well as an increase in T-cell exhaustion^30, 31^. Our data are in keeping with these reports, as we demonstrate a reduction in the estimated proportion of naïve T-cells as well as an increase in memory T-cells with age, several were associated with T-cell exhaustion and/or cellular differentiation. As expected, age related DNA methylation changes were most apparent in datasets from patient cohorts that covered a wide age range^11, 12^. Similarly, in contrast to adult CD patient derived CD8^+^ T-cells, we did not observe major variation in gene transcription of T-cell exhaustion related genes within the pediatric dataset. We therefore speculate that our findings indicate a relative absence of CD8^+^ T-cell exhaustion in children, perhaps contributing to the commonly seen severe and extensive phenotype in childhood onset IBD. Further studies are required to address this concept in more detail. A limitation of our study was that we were unable to combine adult and pediatric datasets due to differences in array types, as well as the lack of matching, publicly available, clinical metadata. Hence, observed age-related differences in CD8^+^ T-cell molecular profiles require confirmation in future studies, ideally containing IBD patients of all ages.

## Conclusions

In summary, our study reveals distinct differences in CD8^+^ T-cell transcriptomes between adult and childhood onset IBD, with the latter lacking correlation with disease outcome. The observed correlation between age and CD8^+^ T-cell gene transcription as well as DNA methylation suggests that our findings may reflect distinct differences in disease pathophysiology between adult and childhood onset IBD. Importantly, our results highlight the need to validate adult clinical biomarkers in independent pediatric patient cohorts.

## Abbreviations

AAV: Anca Associated Vasculitis
CD: Crohn’s Disease
CpG: Cytosine – phosphate - Guanine
DEG: differentially expressed gene
DNAm: DNA methylation
IBD: Inflammatory Bowel Disease
MACS: Magnetic Activated Cell Sorting
PBMC: Peripheral Blood Mononuclear Cells
PCDAI: Pediatric Crohn’s Disease Activity Index
PDCD1: Programmed Cell Death 1
PUCAI: Pediatric Ulcerative Colitis Activity Index
PC: Principle Component
SLE: Systemic Lupus Erythematosus
UC: Ulcerative Colitis
WGCNA: weighted gene co-expression network analysis

