## Supplementary_Methods for "CD8^+^ T-cell transcription and DNA methylation show age specific differences and lack correlation with clinical outcome in pediatric Inflammatory Bowel Disease"

(Supplementary Table 1). In order to account for the fact that disease outcome is not restricted to a single measure, we designed a severity score which considered the number of treatment escalations, escalation to treatment with biologics, peri-anal disease and surgery, as well as unplanned/urgent inpatient admissions (Supplementary Table 2). The score was calculated at 18 months from diagnosis and patients categorised into mild, moderate and severe. All patients were treated according to a standard step-up protocol and treatment decisions made by a multi-disciplinary team making treatment courses of patients highly comparable. Only patients with perineal and/or peri-anal disease received biologics at the point of diagnosis. These were therefore excluded from any analyses aimed at correlating molecular signatures with disease outcome.

### Blood sampling and processing

A peripheral blood sample (volume varied according to age from 10 to 25 ml) for the purification of $CD8^+$ T-cells was taken on the same day as diagnostic endoscopy was performed. All children were treatment naïve at this time point. Additional longitudinal samples (n=62) were taken from a subset of children at 3-month intervals post diagnosis. Disease status was recorded as either 'in remission' or 'active disease' based on their clinical and biochemical disease activity scores (PCDAI and PUCAI) and the treatment history recorded (Supplementary Table 1). All samples were processed immediately and $CD8^+$ T-cells isolated using magnetic bead sorting as detailed below.

### Magnetic bead sorting

A peripheral blood sample of 10 ml (age 4 – 10 years) or 25 ml (age 10 – 17 years) was obtained and $CD8^+$ T-cells extracted using magnetic bead sorting. Briefly, blood was diluted at a 1:2 ratio with MACS rinsing buffer. Peripheral blood mononuclear cells (PBMCs) were isolated by density centrifugation over Ficoll (Histopaque 1077). Following the removal of the plasma, the PBMC interface was transferred to a fresh Falcon tube and $CD8^+$ T-cells were separated by magnetic cell sorting using anti CD8 microbeads (Miltenyi Biotech) as described by the manufacturer. Separation was performed on an AutoMACs Pro Separator (Miltenyi Biotech). Cell purity was assessed regularly on a subset of random samples using Flow cytometry as described below. With an average of 84 % (72.5 - 93.8 %), purity of our samples was found to be similar or higher compared to other published datasets.

### Flow cytometry and cell purity assessment

Flow cytometry was performed on a subset of samples pre- and post-magnetic cell sorting in order to assess purity using FACSCalibur (BD Biosciences). Briefly, following $CD8^+$ T-cell isolation, CD3-PE and CD8-APC antibodies were used (BD Pharmingen), along with the Zombie Aqua Fixable Viability Kit (Biolegend). A total of 8 randomly selected samples were subjected to flow cytometry analysis, and the mean cell purity for $CD8^+$ T-cells was 84 % (72.5 - 93.8 %). Furthermore, we assessed gene expression of lymphocyte subset markers (e.g. CD8, CD14, CD19, CD16) using expression array data, and estimated cellular composition using DNA methylation data. Comparison between our data and published adult datasets confirmed at least similar if not higher purity of our samples (data not shown).

### DNA and RNA extraction

DNA and RNA were extracted simultaneously from isolated $CD8^+$ T-cell samples using AllPrep MiniKit (Qiagen), according to the manufacturer's instructions. DNA and RNA quality were assessed using an Agilent Bioanalyser 2100 and quantified by spectrophotometry using a NanoDrop ND-1000 spectrophotometer. DNA was bisulfite- converted using Zymo DNA methylation Gold kit (Zymo Research).

### Genome wide transcriptional and DNA methylation profiling

Whole genome transcript analysis was performed on 200 ng of total RNA using the Affymetrix Human Gene ST version 2.0 Array (Affymetrix). Genome-wide DNA methylation was profiled using the Illumina EPIC platform (Illumina, Cambridge, UK). All microarray data have been deposited in ArrayExpress, accession numbers: E-MTAB-7923 (expression data) and XXXX (methylation data).

### Publicly available datasets

Gene expression microarray data (Affymetrix Human Gene ST v1.0 and 1.1) from a previously published adult patient cohort diagnosed with CD (n=35) and UC (n=32) was downloaded from ArrayExpress (E-MTAB-331)[4]. Clinical phenotype data available for this cohort included age, gender and disease severity classed as either severe (termed IBD1) or mild (termed IBD2). Secondly, we obtained publicly available CD8$^+$ T-cell derived genome wide DNA methylation profiles (450K Illumina DNA methylation arrays) from two previously published adult patient cohorts. 'Adult cohort 1' (GSE87640)[5], contained a total of 56 CD8$^+$ T-cell samples; 18 CD, 19 UC and 19 healthy control individuals (age range 18 to 63 years). 'Adult cohort 2' included samples obtained from a younger (age 22–34, n=50) and older (age 73–84, n=50) group of healthy adults (GSE59065)[6].

### Bioinformatic analyses

#### *Affymetrix gene expression arrays*

##### Data pre-processing and quality control

The raw signal intensity data were normalized using the variance stabilization and calibration with robust multi-array average (VSNRMA) method using *affy v1.56.0*[7] and quality control was performed using *arrayQualityMetrics v3.34.0*[8]. Samples failing quality control were removed and batch correction was performed using "ComBat" (part of *sva v3.26.0*)[9]. In order to address the possible effect of batch correction on removing transcriptional signatures, the presence of the previously published adult IBD prognostic signature was tested in individual batches and confirmed absence of reproducible/significant clustering. Data was annotated using the *hugene20sttranscriptcluster.db* annotation package. A total of 67 CD, 40 UC, 19 control and 62 follow-up pediatric patient samples and a total of 35 CD adult patient samples were retained for downstream analysis.

**Variance decomposition analysis**

Following quality control, in order to explore the main factors associated with variation within the dataset, principal component (PC) analysis was performed on the normalised, batch corrected pediatric dataset, and the first 10 PCs were examined for correlation with the sample clinical data using Kendall's test statistic for continuous variables and ANOVA for categorical variables (Figure 1A). In addition, we investigated the distribution of sample variation with clinical phenotype more closely by plotting principal components (Figures 1B & C, Supplementary Figures 1A & B).

**Reproduction of the adult $CD8^+$ T-cell prognostic transcriptional signature**

In order to reproduce the previously reported adult $CD8^+$ T-cell prognostic transcriptional signature described by Lee *et al.*[4], we followed the protocol as described in their supplementary methods. Briefly, we used consensus clustering (merging results from k-means and hierarchical clustering from 5000 iterations with an 80% subsampling ratio) to identify optimal clustering within the published dataset described above (*clusterCons v1.0*). Differential expression analysis was then performed on the 2 groups of samples identified (*limma v3.38.3*) and a prognostic signature determined from probes significantly differentially expressed after stringent correction for multiple testing ($p < 0.05$, Holm).

**Application of adult $CD8^+$ T-cell prognostic and T-cell exhaustion signature**

Lists of probes corresponding to the reproduced adult $CD8^+$ T-cell prognostic signature[4] and the exhaustion signature described by Wherry *et al.*[10] (Supplementary Table 4) were used to extract data subsets. Hierarchical clustering was then performed using a Pearson correlation distance metric and average linkage analysis in R using *hclust* (part of the core statistics package) and *heatmap3 v1.1.1* (Figures 2A - D).

**Testing for significance of unsupervised clustering results**

In order to assess clustering patterns for significance, we used the Monte Carlo Reference-based Consensus Clustering package (*M3C*)[11], which uses the Relative Cluster Stability Index (RCSI) and

Monte Carlo adjusted p-values to select the optimal number of clusters (k) rejecting the null hypothesis (k = 1).

**Weighted gene co-expression network analysis (WGCNA)**

WGCNA was used to investigate correlations between gene expression profiles and clinical information, including disease outcomes (Supplementary Table 1) using *WGCNA v1.63*[12] as previously described[13]. Samples from patients given biologics at diagnosis were removed prior to analysis (60 CD and 39 UC samples remaining).

In brief, WGCNA uses unsupervised hierarchical clustering of normalized data, to assess pair-wise correlation between gene expression profiles, implementing soft thresholding to assign a connection weight to each gene pair[12], in order to reduce the dimensions of the dataset by grouping highly correlated genes into modules (eigengenes). Eigengene significance (correlation between sample trait and eigengene) and p-values are then calculated for the whole module instead of individual genes, greatly alleviating multiple-testing.

For our analysis, soft thresholding power was chosen based on the scale-free topology model parameters (scale independence and mean connectivity). Gene networks were constructed and modules identified from the resulting topological overlap matrix (dissimilarity correlation threshold = 0.01, minimum module size: n = 30; deepSplit = 2). The resulting modules (eigengenes) were aligned to the clinical information, tested for correlation (using Pearson's correlation) and Student's asymptoptic p-values calculated. False discovery rate (FDR) correction for multiple testing of the resulting p-values was performed using the Benjamini-Hochberg procedure.

***DNA methylation analysis***

**Data pre-processing and quality control**

Illumina EPIC arrays provided quantitative measures of DNA methylation (DNAm) for $CD8^+$ T-cells from 66 patients, at single CpG resolution (>850,000 sites) covering the whole genome. DNAm data was processed using the *minfi* package v1.28[14], specifically the "read.metharray" function, to extract beta values from raw IDAT files. Data was then normalized based on control probes on each array using functional normalization[15]. Starting with a total of 866,238 probes present on the EPIC array, the following probes were filtered out: polymorphic CpG[16], located on sex chromosome, potential to cross hybridize to several regions of the genome[16], poor quality as measured by a detection p value > 0.05 in at least 1 % of samples. This filtering left 792,401 CpGs for analysis. Batch correction was performed using "ComBat" on array ID (part of sva v3.30.0)[9].

#### Correlating DNA methylation with clinical outcome data

To explore the main factors associated with variation in DNAm data, Principal Component (PC) Analysis was performed on the normalized, batch corrected dataset. The first 10 PCs were examined for correlation with clinical data using Spearman's correlation for continuous variables and ANOVA for categorical variables (Figure 5A). In addition, the distribution of PC1 and PC2 was compared with clinical data to explore the major contributors to variation in DNAm (Figure 5B and data not shown).

#### Differential Methylation Analysis

Differential methylation with severity score was tested at each CpG using limma v3.38.3[1]. Models included covariates for age and gender. To be considered significantly differentially methylated, CpGs needed to have an association p-value $<9\times10^{-}8$ (selected according to current recommendations)[17] and an absolute methylation difference between the most severe and least severe of 0.05 (delta beta > 0.05).

Differential methylation with age was tested in Adult cohort 2 (GSE59065), as it had the greatest age range. Published DNAm data was processed as above, using annotation for the Illumina 450K array[18]. Differential methylation at individual CpGs was tested using limma v3.38.3[19], while covarying for gender, using the same threshold criteria as described above[17].

To compare differential DNAm to the known lists of $CD8^+$ T-cell gene expression signatures, CpGs were associated with adjacent genes. A CpG was assigned to a gene based on proximity to a transcript from Ensembl Genes 99, GRCh37.p13, collected from BioMaRt[20]. CpGs were classed as being associated with a gene if they were located between 1,500 bp upstream of the transcription start and 300 bp downstream of the transcript end. CpG to transcript associations were then aggregated by Ensembl Gene stable ID.

**Epigenetic Clock and Cell subset estimation Analysis**

The normalized beta values as provided under GSE87640 (Adult cohort 1) and GSE59065 (Adult cohort 2) were used to estimate epigenetic age. Using the Horvath epigenetic clock[21], with the clock's normalization applied, the DNAm based age of each sample was estimated. As an extension, the estimated abundance levels of $CD8^+$ T naïve and $CD8^+$ T memory cells ($CD8^+CD28^-CD45RA^-$ T-cells) were obtained from the 'Advanced Blood Analysis' of the online DNAm age predictor[21]. These measures indicate ordinal abundances from a regression with flow-sorted counts from other datasets. Hence, resulting data allows comparison of cell-subset proportions across samples over chronological age within a cohort.

A.

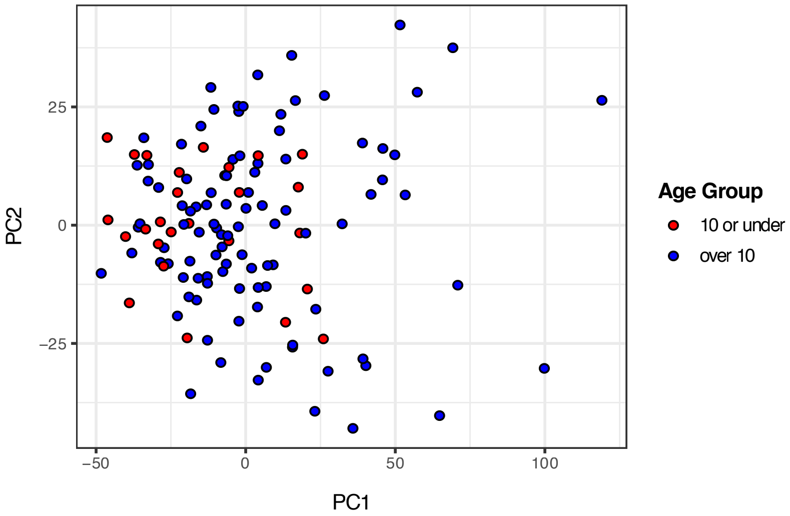

PC2
25
0
−25
−50
0
50
100
PC1
Age Group
10 or under
over 10


B.

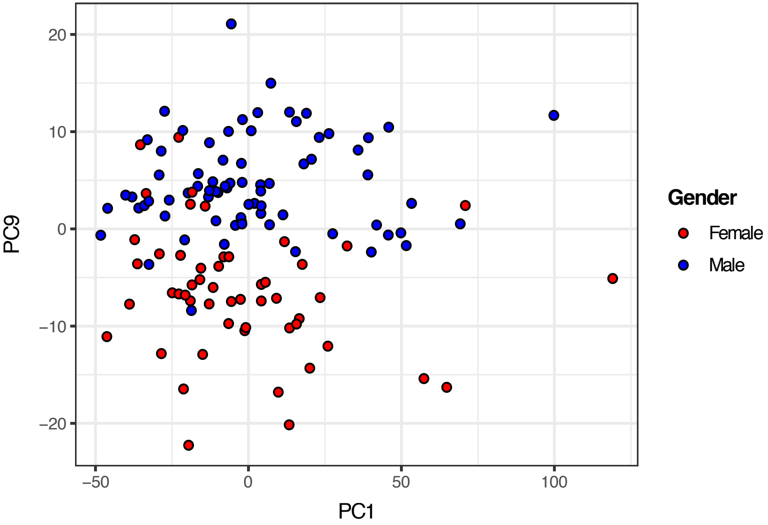

PC9
20
10
0
−10
−20
−50
0
50
100
PC1
Gender
Female
Male

**Supplementary Figure 1**: Genome wide transcriptional profiles of CD8$^{+}$ T-cells obtained from children newly diagnosed with Crohn's Disease (n=67), Ulcerative Colitis (n=40) and healthy controls (n=19). PCA plots were generated following pre-processing and batch correction **A)** PC1 versus PC2 with samples labeled by age: 10 years or under (n=26, red) or over 10 years of age (n=100, blue). **B)** Showing PC1 versus PC9 with samples labelled by gender: male (n=74, blue), female (n=52, red).

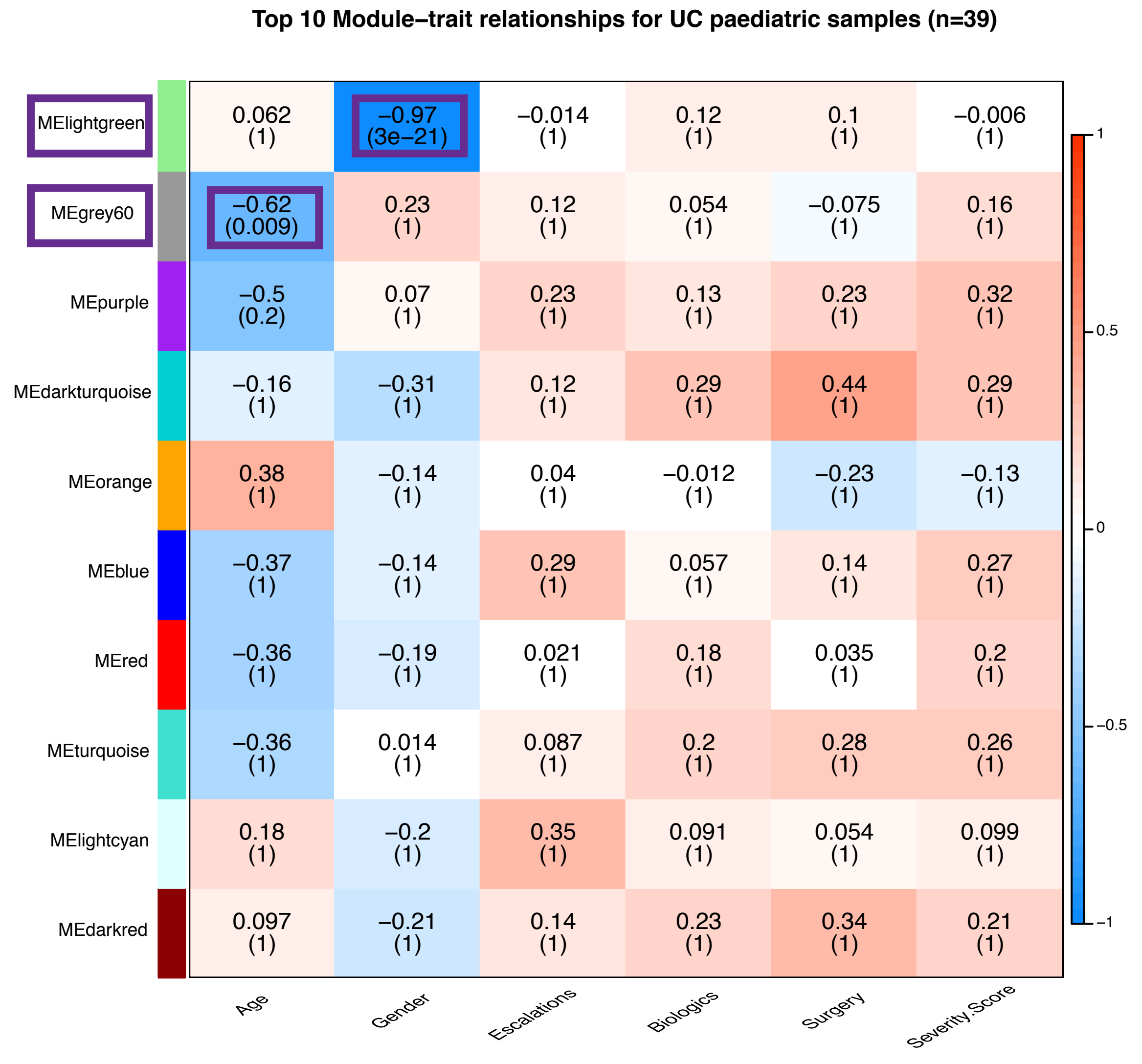
Top 10 Module–trait relationships for UC paediatric samples (n=39)
MElightgreen
MEgrey60
MEpurple
MEdarkturquoise
MEorange
MEblue
MEred
MEturquoise
MElightcyan
MEdarkred
0.062 (1)
−0.97 (3e−21)
−0.014 (1)
0.12 (1)
0.1 (1)
−0.006 (1)
−0.62 (0.009)
0.23 (1)
0.12 (1)
0.054 (1)
−0.075 (1)
0.16 (1)
−0.5 (0.2)
0.07 (1)
0.23 (1)
0.13 (1)
0.23 (1)
0.32 (1)
−0.16 (1)
−0.31 (1)
0.12 (1)
0.29 (1)
0.44 (1)
0.29 (1)
0.38 (1)
−0.14 (1)
0.04 (1)
−0.012 (1)
−0.23 (1)
−0.13 (1)
−0.37 (1)
−0.14 (1)
0.29 (1)
0.057 (1)
0.14 (1)
0.27 (1)
−0.36 (1)
−0.19 (1)
0.021 (1)
0.18 (1)
0.035 (1)
0.2 (1)
−0.36 (1)
0.014 (1)
0.087 (1)
0.2 (1)
0.28 (1)
0.26 (1)
0.18 (1)
−0.2 (1)
0.35 (1)
0.091 (1)
0.054 (1)
0.099 (1)
0.097 (1)
−0.21 (1)
0.14 (1)
0.23 (1)
0.34 (1)
0.21 (1)
Age
Gender
Escalations
Biologics
Surgery
Severity.Score
1
0.5
0
−0.5
−1

**Supplementary Figure 2**: Application of Weighted Gene Co-expression Network Analyses (WGCNA) to CD8$^+$ T-cell transcriptional profiles derived from pediatric patients diagnosed with UC (n=40), showing the top 10 gene modules and their association with clinical phenotype (age at diagnosis, gender, number of treatment escalations, treatment escalation to biologics, surgery and summary severity score). Only correlations with age (grey60) and gender (lightgreen) reached statistical significance ($p<0.05$). Numbers indicate degree of correlation between module and trait (top) and corrected p-value evaluating statistical significance of association (in parentheses).

A.

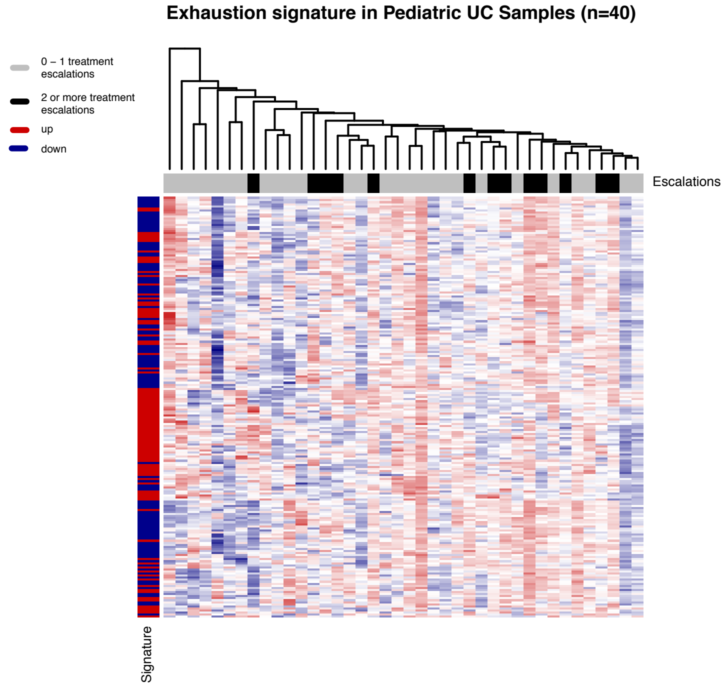
Exhaustion signature in Pediatric UC Samples (n=40)
0 – 1 treatment escalations
2 or more treatment escalations
up
down
Escalations
Signature

B.

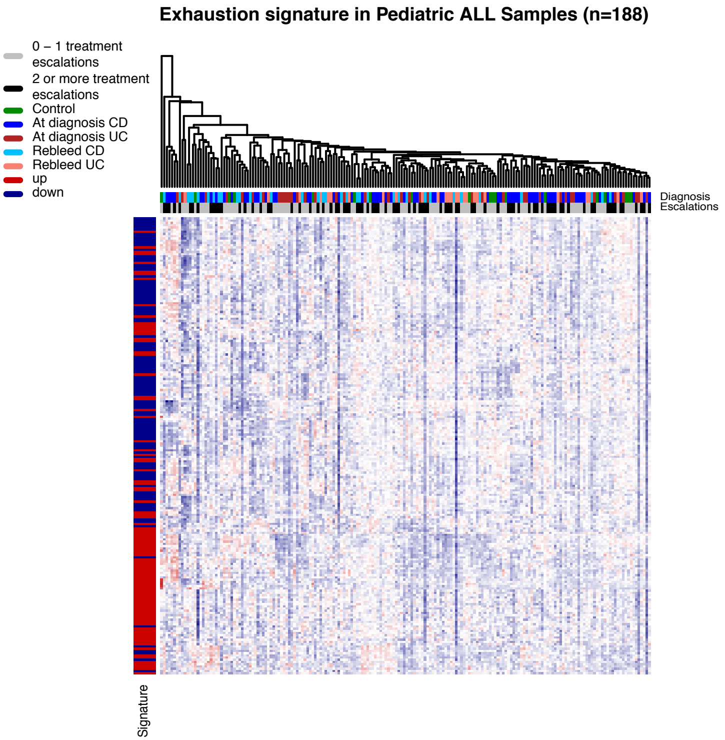
Exhaustion signature in Pediatric ALL Samples (n=188)
0 – 1 treatment escalations
2 or more treatment escalations
Control
At diagnosis CD
At diagnosis UC
Rebleed CD
Rebleed UC
up
down
Diagnosis
Escalations
Signature

**Supplementary Figure 3**: Transcriptional variation of genes associated with T-cell exhaustion in $CD8^+$ T-cells. Genes forming a transcriptional T-cell exhaustion signature (Wherry *et al.*)[24] were selected from genome-wide transcriptomes and subjected to hierarchical clustering. Heatmap and hierarchical clustering trees are displayed for samples obtained from **A**) Children newly diagnosed with UC (n=40) and **B)** all pediatric patient derived $CD8^+$ T-cell transcriptomes (n=188, including controls and longitudinal follow-up samples). No significant clusters were identified and patients with mild (i.e. 0-1 treatment escalations in grey), or moderate to severe (i.e. 2 or more treatment escalations in black) outcome are distributed equally. The y axis indicates expression of genes according to reference signature (red=upregulated, blue=downregulated).

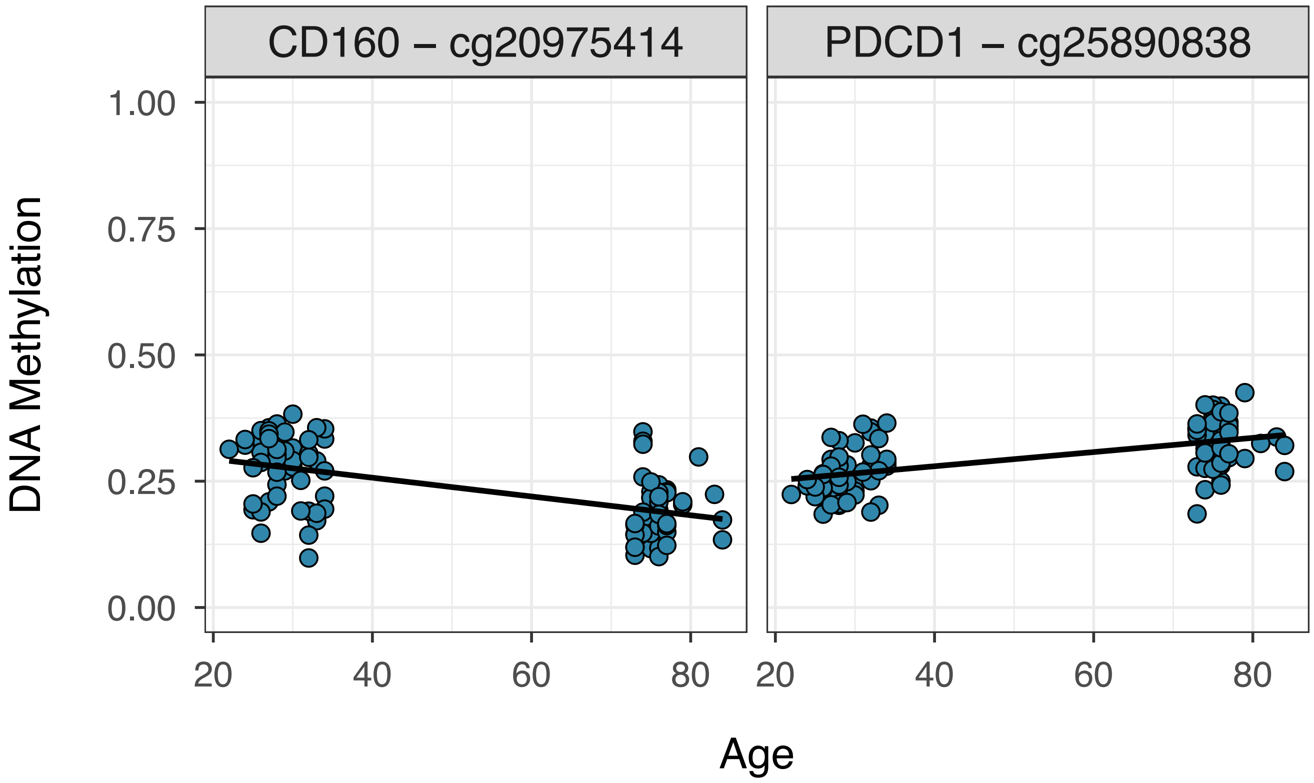
CD160 – cg20975414
PDCD1 – cg25890838
DNA Methylation
1.00
0.75
0.50
0.25
0.00
20
40
60
80
Age

**Supplementary Figure 4**: Age dependent DNA methylation changes in $CD8^+$ T-cells. Differential DNA methylation analysis was performed on a published dataset (Adult Cohort 2, Tserel *et al.*)[12] comparing methylomes derived from young versus old adults. Amongst the significantly differentially methylated CpGs were several associated with T-cell exhaustion. Displayed are two examples: CD160 and Programmed Cell Death 1 (PDCD1). The y-axis indicates level of DNA methylation displayed as beta-value ranging from 0 (unmethylated) to 1 (methylated). The x-axis displays chronological age. Plots are labelled as gene name followed by CpG ID (Illumina DNA methylation array).
