## Supplementary_Table_2 for "CD8^+^ T-cell transcription and DNA methylation show age specific differences and lack correlation with clinical outcome in pediatric Inflammatory Bowel Disease"

### Supplementary Table 2. Disease outcome severity score

| Crohn's disease | Ulcerative colitis |
|---|---|
| **1. Number of treatment escalations** | **1. Number of treatment escalations** |
| 0-1: **0** 2: **1** >=3: **2** | 0-1: **0** 2: **1** >=3: **2** |
| **2. Biologics** | **2. Biologics** |
| No: **0** Yes: **2** | No: **0** Yes: **2** |
| **3. Crohn's related surgery** | **3. UC related surgery (colectomy)** |
| No: **0** Yes: **2** | No: **0** Yes: **2** |
| **4. Perianal disease** | **4. Steroid-free remission at 3 months from diagnosis (PUCAI score < 10)** |
| Absent: **0**<br>Medical management: **1**<br>Surgical management: **2** | Yes: **0** No: **2** |
| **5. Unplanned/urgent inpatient days** | **5. Unplanned/urgent inpatient days** |
| 0-2: **0** 3-4: **1** >=5: **2** | 0-2: **0** 3-4: **1** >=5: **2** |
| **Outcome based on total score**<br>**0-1: Mild**<br>**2-4: Moderate**<br>**5-10: Severe** | **Outcome based on total score**<br>**0-1: Mild**<br>**2-4: Moderate**<br>**5-10: Severe** |
